# The SONATA Data Format for Efficient Description of Large-Scale Network Models

**DOI:** 10.1101/625491

**Authors:** Kael Dai, Juan Hernando, Yazan N. Billeh, Sergey L. Gratiy, Judit Planas, Andrew P. Davison, Salvador Dura-Bernal, Padraig Gleeson, Adrien Devresse, Benjamin K. Dichter, Michael Gevaert, James G. King, Werner A. H. Van Geit, Arseny V. Povolotsky, Eilif Muller, Jean-Denis Courcol, Anton Arkhipov

**Affiliations:** Allen Institute for Brain Science, Seattle, WA, USA; Blue Brain Project, École Polytechnique Fédérale de Lausanne, Lausanne, Switzerland; Paris-Saclay Institute of Neuroscience UMR 9197, Centre National de la Recherche Scientifique/Université Paris Sud, France; State University of New York Downstate Medical Center, Brooklyn, NY, USA; Nathan Kline Institute for Psychiatric Research, Orangeburg, NY, USA; Department of Neuroscience, Physiology and Pharmacology, University College London, UK; Department of Neurosurgery, Stanford University, Stanford, CA, USA; Biological Systems and Engineering, Lawrence Berkeley National Laboratory, Berkeley, CA, USA

## Abstract

Increasing availability of comprehensive experimental datasets and of high-performance computing resources are driving rapid growth in scale, complexity, and biological realism of computational models in neuroscience. To support construction and simulation, as well as sharing of such large-scale models, a broadly applicable, flexible, and high-performance data format is necessary. To address this need, we have developed the Scalable Open Network Architecture TemplAte (SONATA) data format. It is designed for memory and computational efficiency and works across multiple platforms. The format represents neuronal circuits and simulation inputs and outputs via standardized files and provides much flexibility for adding new conventions or extensions. SONATA is used in multiple modeling and visualization tools, and we also provide reference Application Programming Interfaces and model examples to catalyze further adoption. SONATA format is free and open for the community to use and build upon with the goal of enabling efficient model building, sharing, and reproducibility.

## Introduction

Modern systems neuroscience faces ever-widening streams of data on composition, connectivity, and *in vivo* activity of brain networks (e.g., (Gouwens et al., 2018a; Jiang et al., 2015; Kasthuri et al., 2015; Lee et al., 2016; Markov et al., 2012; Oh et al., 2014; Tasic et al., 2018; de Vries et al., 2018)), supported by major funding initiatives around the world (Amunts et al., 2016; Bouchard et al., 2016; Hawrylycz et al., 2016; Koch and Jones, 2016; Martin and Chun, 2016; Vogelstein et al., 2016). Turning these complex data into knowledge is a challenging task requiring systematic analysis and modeling. Detailed, data-driven modeling in particular will be essential to integrate the experimentally observed hundreds of cell types, intricate connectivity rules, and complex patterns of neuronal dynamics into predictive computational frameworks (Einevoll et al., 2019).

For this task, scientists need tools that are up to the challenge. Simulation engines, such as NEURON (Carnevale and Hines, 2006), NEST (Gewaltig and Diesmann, 2007), Brian (Goodman and Brette, 2008), GENESIS (Bower and Beeman, 1997), MOOSE (Ray and Bhalla, 2008), Xolotl (Gorur-Shandilya et al., 2018), and others offer high computational performance, and recently a number of software interfaces (e.g., neuroConstruct (Gleeson et al., 2007), PyNN (Davison et al., 2009), NetPyNE (Dura-Bernal et al., 2019), Open Source Brain (Gleeson et al., 2018), and the Allen Institute’s Brain Modeling ToolKit (BMTK, https://alleninstitute.github.io/bmtk/; (Gratiy et al., 2018)) have been developed that allow users to interact with these engines without mastering the underlying programming environments. However, the utility of these tools is limited without a broadly applicable, flexible, and high-performance modeling data format. The current evolution of typical workstyles towards collaborative team projects demands standardized formats for model sharing and reproducibility, as well as for interoperability between tools. Meanwhile, high computational performance of such formats becomes increasingly important to enable efficient representation of growing biological complexity of models.

While existing solutions, such as the XML-based data format NeuroML (Cannon et al., 2014; Gleeson et al., 2010), the PyNN language (Davison et al., 2009), and the NSDF standard for simulator output (Ray et al., 2016), have proven useful, major challenges remain and are felt acutely in the case of large data-driven models. One problem is a performance bottleneck: storing data about thousands of neurons or millions of synapses in verbose text-based files produces a large disk space footprint and may be challenging for reading/writing in parallel compute environments. Another is that existing formats describe either static models or simulation outputs, but not both. And, for broad adoption of a modeling data format, it needs to be flexible enough to represent a variety of model types (point neuron, biophysically detailed, etc.) and compatible with more specialized formats (e.g., SWC for neuronal morphologies (Cannon et al., 1998)), without compromising computational performance.

Notably, similar challenges exist in experimental neuroscience (see, e.g., (Koch and Reid, 2012)). The situation is improving due to initiatives for experimental data formats, such as NWB:N (Ruebel et al., 2019), BIDS (Gorgolewski et al., 2016), Loom (https://linnarssonlab.org/loompy), or spacetx-starfish (https://github.com/spacetx/starfish), but for many types of experimental data the community is still far from a widespread adoption of universally agreed-upon formats. These challenges contribute to difficulties in closing the virtuous experiment/modeling loop and to the overall reproducibility crisis (Baker, 2016; Goodman et al., 2016; Koch and Jones, 2016)).

Here we present the SONATA (Scalable Open Network Architecture TemplAte) data format, which provides an open-source framework for representing neuronal circuits, simulation configurations, and simulation outputs. The format has been jointly developed by the Allen Institute and the Blue Brain Project to facilitate exchange of their large scale cortical models (e.g., (Arkhipov et al., 2018; Billeh et al., 2019; Markram et al., 2015)) and is supported by these organizations’ software tools, such as BMTK (https://alleninstitute.github.io/bmtk/; (Gratiy et al., 2018)). Support for the format has also been added by other simulation tools -- pyNeuroML (Cannon et al., 2014; Gleeson et al., 2010), PyNN (Davison et al. 2009), and NetPyNE (Dura-Bernal et al., 2019) -- and an interface between SONATA and the NWB:N format (Ruebel et al., 2019) for neurophysiological data has been developed.

As described below, SONATA utilizes computationally efficient binary formats for storing large datasets while also offering text-based formats for easy editing of less data-rich model components. SONATA represents all aspects of models and simulations, from network structure, to simulation parameters, to input and output activity. It provides much flexibility for describing models at different levels of resolution, including hybrid models. Importantly, because SONATA is already supported by a number of widely used tools and applications, users can get all of the benefits of the format with no extra work on their part. Full specification of the format can be found at the SONATA GitHub page (https://github.com/AllenInstitute/sonata), along with the open-source reference application programming interfaces (APIs). To enable broad applications in the field, SONATA is freely available and open to the community.

## Results

### Overview of the SONATA format

The major object in SONATA is the model network (**Fig. 1**), which consists of **nodes** of two types: explicitly simulated nodes and virtual nodes (the latter only providing inputs to the simulated system). In both cases, nodes are grouped in one or more **populations** for convenience. Nodes within and between populations are connected via **edges**. Simulations of model networks are performed by applications that load SONATA files. Locations of these files and also parameters of simulation (e.g., the time step and temperature) are stored in the SONATA configuration (**“config”**) files. Finally, SONATA also provides specifications to store the incoming activity or simulation output, in the form of events (spikes) or time series.

**Figure 1.**
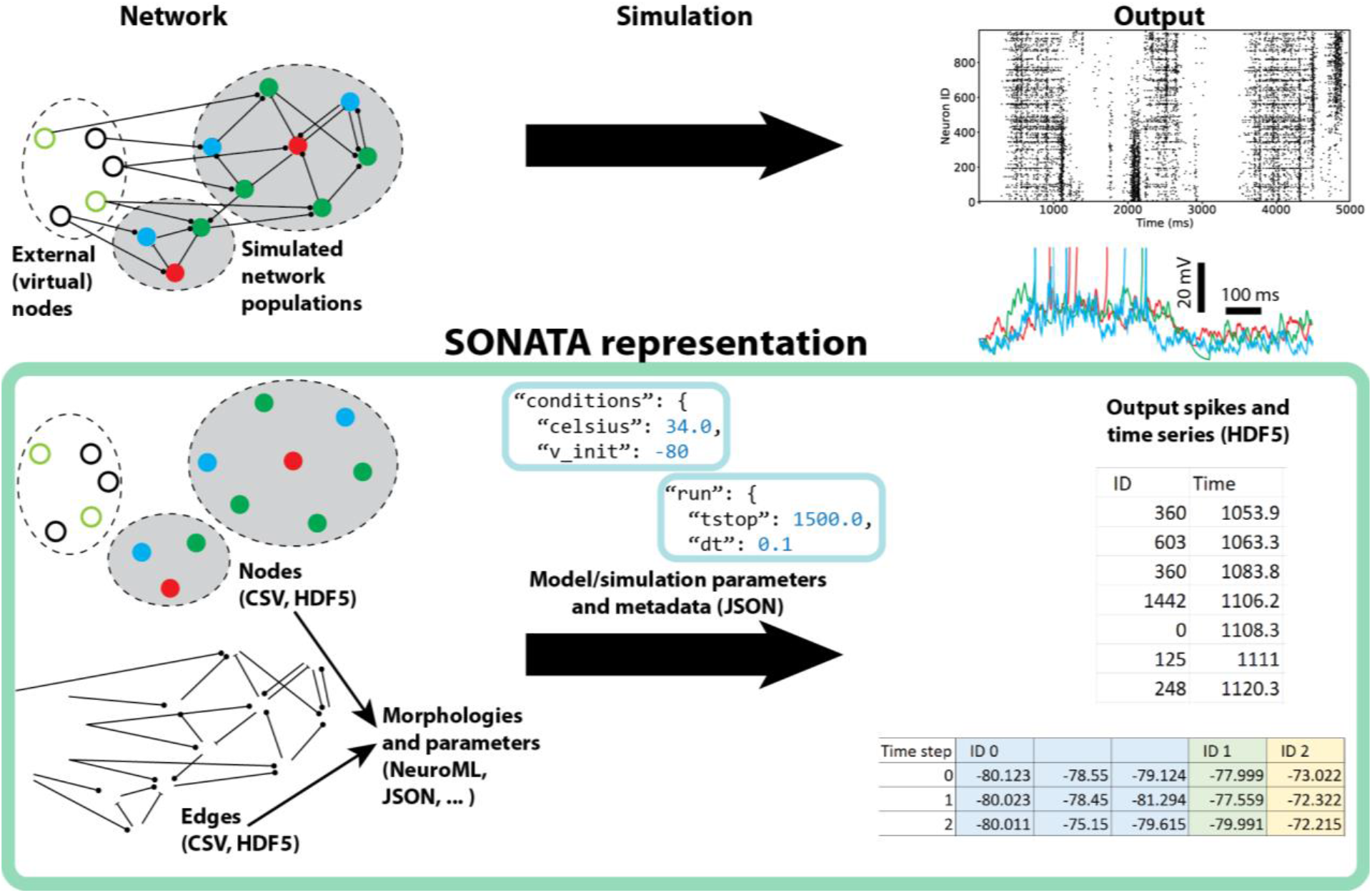
Overview of the SONATA data format. (Top) A simulated model consists of one or more explicitly simulated network populations and external sources (virtual nodes) that provide inputs into the simulated populations. During and after simulation, output is created characterizing dynamics in the simulated model. (Bottom) The SONATA data format reflects the major components of simulation in dedicated file structures. Information about the model is stored in files (CSV and HDF5) describing nodes and edges of the network (left). Model metadata (e.g., path relations between files on disk) and information about simulation are stored in JSON configuration files (middle). The spiking and time series output is stored in a tabular format, taking advantage of the HDF5 technology (right). In the case of time series (bottom right), multiple variables can be stored for individual nodes (in this example, node ID 0 has three variables stored), which can correspond, e.g., to multiple compartments of a neuron.

SONATA relies on existing file formats, HDF5, CSV, and JSON (see Methods), which ensures that files can be read/written by existing libraries and applications and used on all major operating systems. The SONATA specification on top of these formats accommodates multiple cell and synapse model types and is designed to optimally handle a heterogeneous network. To achieve flexibility in defining models, SONATA provides recipes for storing arbitrary attributes, with some attribute names being reserved for basic standardization.

Below, we describe the details of these elements of the SONATA format. A more complete description is given in the **Online Documentation** (https://github.com/AllenInstitute/sonata/blob/master/docs/SONATA_DEVELOPER_GUIDE.md).

### Node and edge types

Both nodes and edges can have **attributes** describing biological details (e.g. cell or synapse properties). One major benefit of the SONATA format is its flexibility: while a small number of attributes are reserved, users can create their own attributes for nodes or edges. Furthermore, attributes can be described either individually for each node or more globally for whole subsets of nodes (same for edges), due to the concepts of **node types** and **edge types**. It is up to the user to decide which attributes are stored on a per-type basis and which should be stored individually for each node or edge. Since the number of node/edge types in a network model is usually much smaller than the number of nodes or edges, the node/edge type files are stored in the plain-text CSV tabular format. This makes it easy for modelers to change and update the network *en-masse* through a text editor. For example, **Table 1** shows five different node types, three of which (node_type_id 100, 101, and 102) are biophysically detailed models and two (node_type_id 103 and 104) are much simpler, point neuron models. Whereas the total number of nodes in this network can be many thousands, the five entries in **Table 1** succinctly describe many attributes of the nodes.

**Table 1:**
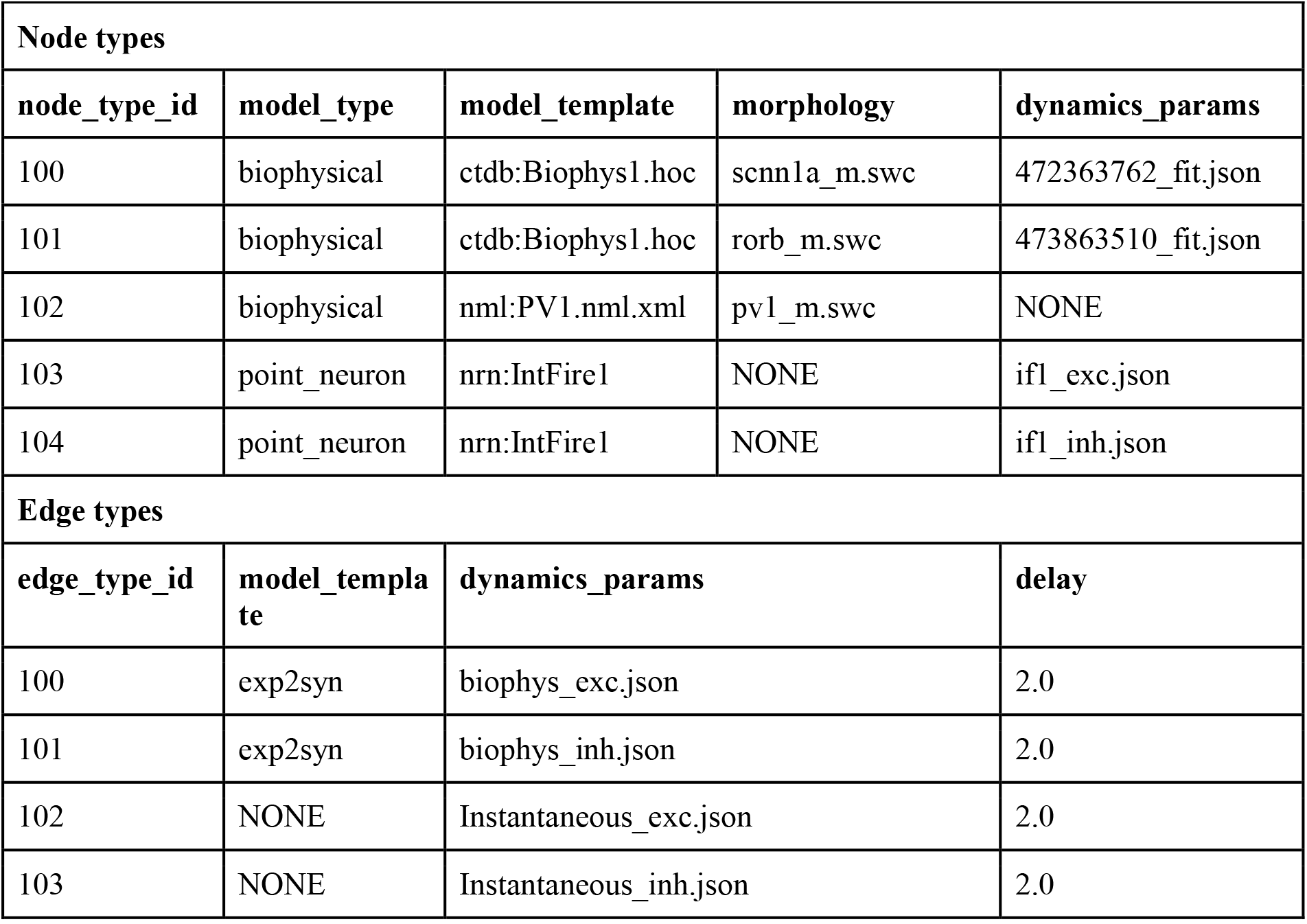
Examples of “node types” and “edge types”. In a network model, all individual nodes belonging to a particular node type share the respective attributes, and likewise all edges belonging to the same edge type share attributes of that type.

The lists of attributes and instructions for constructing individual nodes are determined by each node type’s “model_type” (**Table 1**). The reserved values are “biophysical”, “single_compartment”, “point_neuron”, or “virtual”. The mechanisms required for cell models are described by “model_template”, with possible values including references to a NeuroML2 file or a NEURON hoc template. The reserved “morphology” attribute references a morphology file (e.g., in the widely used SWC format) and the “dynamics_params” references files that can be optionally used to initialize or overwrite electrophysiological attribute values defined by the template. In **Table 1**, node types 100 and 101 are built using hoc templates from the Allen Cell Types Database (http://celltypes.brain-map.org), which take parameter values form the JSON files in “dynamics_params”. Node type 102 uses a NeuroML template file; dynamics_params = NONE means that default values from the NeuroML model_template are used. Node types 103 and 104 are NEURON built-in IntFire1 point processes taking parameter values from the JSON files under “dynamics_params”.

Edge types are described in similar ways (**Table 1**). The “model_template” attribute determines the synaptic model via a template file or a synaptic type defined in a particular simulator, e.g., NEURON’s exp2syn, whereas the optional “dynamics_params” initializes or overwrites the parameters of the synaptic mechanisms, e.g., time of rise and decay of synaptic conductance. Other reserved attributes include synaptic weight, delay, and the afferent and efferent locations of synapses (only the delays are shown in **Table 1**).

### Nodes

Individual attributes of nodes are listed in “node tables”, stored as HDF5 files. As discussed, users decide which attributes to store in node-type CSV and which in node table HDF5. For example (**Fig. 2A**), one can store only the coordinates of neurons (x, y, z locations) in the node table with a pointer (the *node_type_id*) to the node types table for repeated information such as morphology (see example in **Table 1**). Alternatively, each neuron may have its own unique morphology (**Fig. 2B**), and in that case the node table contains both the coordinates and the morphology attribute.

**Figure 2.**
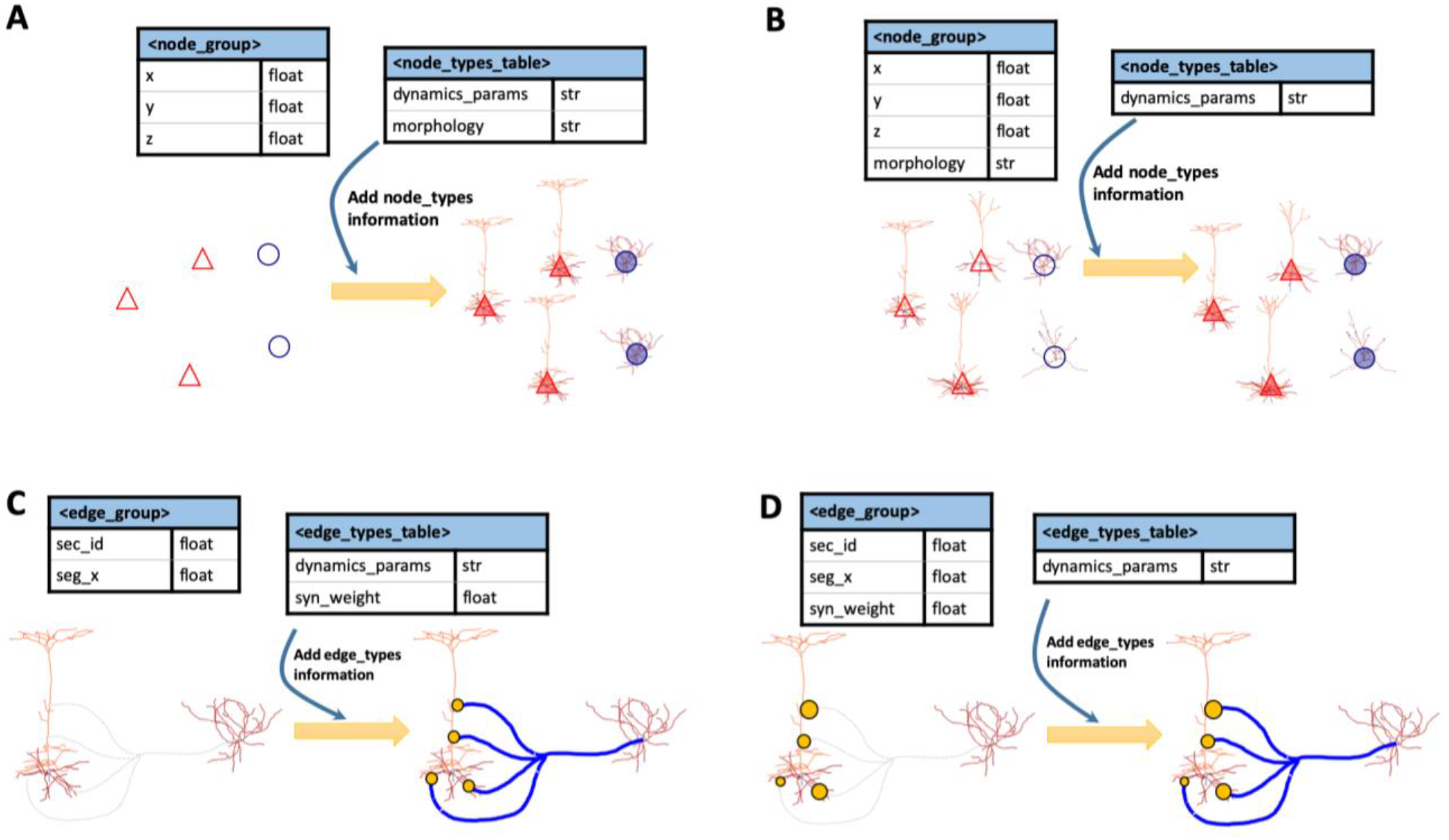
Nodes and edges in SONATA format. (A, B) Two examples are shown that demonstrate how for each node one can find its model attributes in either the node_group (for individually unique attributes) or the node_types table (for globally shared attributes). In (A), the unique attributes are only the node locations (x, y, z), indicated by empty triangles and circles on the left. Morphology and dynamic parameters are shared among multiple nodes within a type. Hence, all red triangles share the same morphology, as do blue circles (right). In (B), the morphology is unique for each node. The dynamics_params is the only attribute specified at the type level; it is assigned to each node, as indicated by the triangles and circles being filled with color on the right. (C, D) Same for edges. In (C), the synapse locations are stored individually for each edge, whereas synaptic weights and dynamics_params attributes are stored at the edge type level, as indicated by the uniform circle size and colored connections on the right. (“dynamics_params” attributes here determine the dynamical properties of synapses, such as the time of rise and time of decay of synaptic conductance). (D) The synapse locations as well as synaptic weights are stored individually (hence different circle sizes), whereas the dynamics_params attributes are stored at the edge type level.

SONATA allows for nodes to be hierarchically organized into **populations** and **groups**. Different populations may be stored in different files, allowing modelers to mix and reuse populations between simulations. For example, one may study one brain region -- say, visual area V1 -- in one simulation and visual area V2 in another simulation, and then build a simulation of V1 and V2 together using the two populations without the need to create new nodes files. Within a population, there is one or more node groups, each group using a homogeneous collection of node attributes. This is useful for hybrid simulations. For example, compartmental neuron models often have many more (and radically different) attributes than integrate-and-fire models. Thus, for mixed populations it is practical to store attributes of compartmental and integrate-and-fire nodes in different groups. Note that nodes of multiple types may be stored in each group, as long as all the nodes in the group have the same lists of attributes. The SONATA implementation of populations and groups utilizes HDF5 groups and datasets (see **Online Documentation**).

### Edges

An edge typically represents a synapse from one neuron to another. Like for nodes, shared attributes of many edges can be stored in CSV edge type files and individual attributes in HDF5 edge tables files (**Fig. 2C, D**). Users decide which attributes belong to edge types and which to edge tables. In the edge tables, edges are grouped together into **edge populations**. Each edge population contains directed connections between nodes in one node population to nodes in another population (the target and source populations can be the same). Each edge identifies the node_id of the source node and the node_id of the target. There may be multiple edges for a single source/target pair. As with nodes, each edge population consists of one or more edge groups. One edge group contains edges with the same list of attributes.

Continuing our example of a model of V1 and V2 above, one can use one edge population for all connections from V1 to V2, another for V2 to V1, another for V1 to V1, and one more for V2 to V2. The specific partition is again up to users, but has to be consistent with the partition of nodes into populations. Within the V1-to-V1 edge population, one may need to have two edge groups. One edge group would be used for connections to biophysically detailed cell models, containing, for example, attributes of synapse location on the dendritic tree of the target cell, local synapse strength, time delay specific to that particular edge, and many others. The other edge group would be used for connections to point-neuron models, perhaps containing only the synaptic weight.

For technical details and benchmark examples of SONATA representation of edges, see Methods.

### Simulation configuration

SONATA provides a framework for storing the information about the location of the files describing the model, as well as parameters of the simulation and metadata. This information is stored in the **config** files that tie all the network, circuit, and output components together (**Fig. 1).** The SONATA configuration files, the **primary config**, the **circuit config**, and the **simulation config**, are JSON files containing key/value pairs. **Table 2** lists the keys required in each of these files (see **Online Documentation** for details).

**Table 2.**
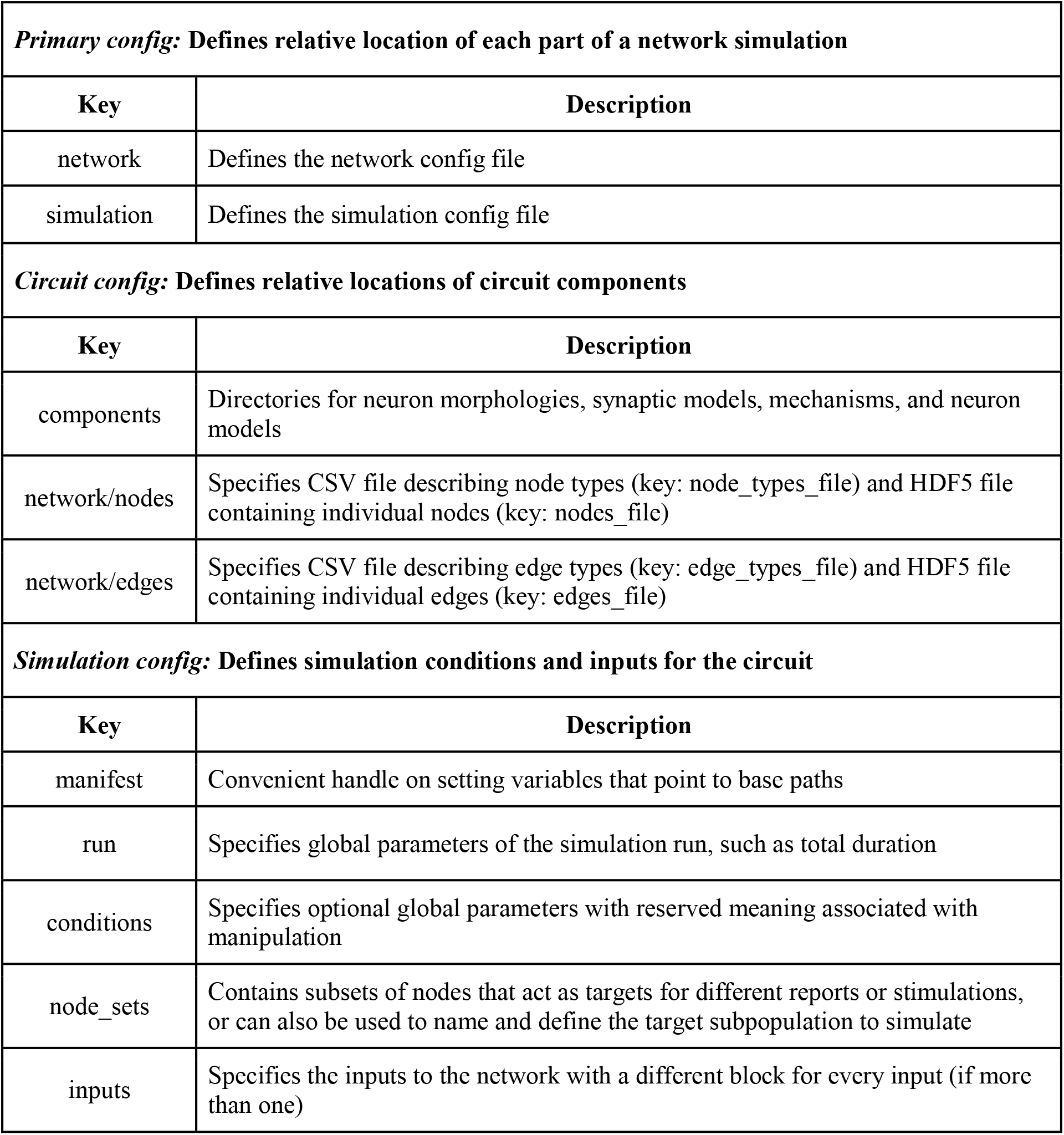
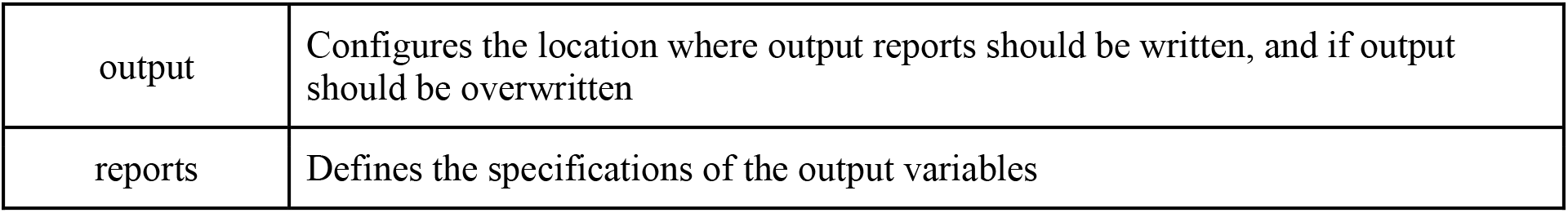
Summary of the *config* files. Representative components are listed; additional entries can be used as described in the **Online Documentation**.

| **Primary config: Defines relative location of each part of a network simulation** |  |
| --- | --- |
| **Key** | **Description** |
| network | Defines the network config file |
| simulation | Defines the simulation config file |
| **Circuit config: Defines relative locations of circuit components** |  |
| **Key** | **Description** |
| components | Directories for neuron morphologies, synaptic models, mechanisms, and neuron models |
| network/nodes | Specifies CSV file describing node types (key: node_types_file) and HDF5 file containing individual nodes (key: nodes_file) |
| network/edges | Specifies CSV file describing edge types (key: edge_types_file) and HDF5 file containing individual edges (key: edges_file) |
| **Simulation config: Defines simulation conditions and inputs for the circuit** |  |
| **Key** | **Description** |
| manifest | Convenient handle on setting variables that point to base paths |
| run | Specifies global parameters of the simulation run, such as total duration |
| conditions | Specifies optional global parameters with reserved meaning associated with manipulation |
| node_sets | Contains subsets of nodes that act as targets for different reports or stimulations, or can also be used to name and define the target subpopulation to simulate |
| inputs | Specifies the inputs to the network with a different block for every input (if more than one) |
| output | Configures the location where output reports should be written, and if output should be overwritten |
| reports | Defines the specifications of the output variables |

The circuit config contains pointers to the files with the information about nodes and edges that describe the network being simulated. The simulation config describes properties unique to a specific simulation run, such as the inputs the network receives, the simulation parameters (for example, duration, time-step), optional parameters such as the temperature, the outputs to be recorded (for instance spike times, membrane potentials, internal calcium concentrations, etc.), paths to writing the outputs, and others. Both the simulation config and the circuit config may contain a manifest block that defines the paths to be used/reused throughout the JSON file. The primary config simply points to the simulation and circuit configs.

Separating of config files in this manner provides flexibility to mix and match models and simulations. For example, one can use a single circuit config and multiple simulation configs to run many simulations of one model under different conditions, or alternatively use multiple circuit configs with one simulation config to study multiple circuits under identical conditions.

### Input and output activity

In addition to representing models, SONATA also describes dynamical variables such as spikes and time series, which is necessary for representing incoming activity or output of simulations. For these types of data, SONATA’s format is in many ways similar and consistent to the experimental neurophysiology format NWB:N (Ruebel et al., 2019), the two formats having been developed approximately simultaneously and with mutual influences due to interactions between the two developer communities. Both are designed to be optimal for large-scale recordings or simulations. At present, the SONATA output format and NWB:N are maintained in separate projects, but conversion between the two is straightforward and is achieved by a tool described below (see **Ecosystem support**). In the future, it may be desirable to achieve full integration between NWB:N and SONATA.

#### Activity format design

The SONATA activity format (also referred to as **reports**) is designed to efficiently support three types of data: spike trains, time series for node elements (e.g., membrane voltage or Ca^2+^ concentration in cell compartments) and time series that are not associated with specific node elements (such as voltages recorded with extracellular probes). The file formats are based on HDF5.

The data stored in a spike train report consists of a series of node identifiers and spike times, stored in separate HDF5 datasets. For maximum flexibility, the standard allows the datasets to be sorted according to three different criteria: by node ID, by spike time, or unsorted.

A node element report consists of a set of variables which are sampled at a fixed rate for some elements of interest from a selected set of cells. Typically, the elements are electrical compartments, but other elements can be used as well, such as individual synapses. The time series associated with each element can be membrane voltage, synaptic current, or any other variable. In the report, a simulation **frame** is the set of all values reported at a given timestamp and a **trace** is the full time series of all values associated with one element (**Fig. 3A**). The requirements we followed in designing the node element report were: (i) support for large data sets both in total size (terabytes) and number of elements (millions of cells using multi-compartment models), (ii) random read access to specific frames and elements within a frame, (iii) high performance for different read access patterns (especially full frames and full cell traces) and (iv) high performance sequential parallel writing of full frames.

**Figure 3.**
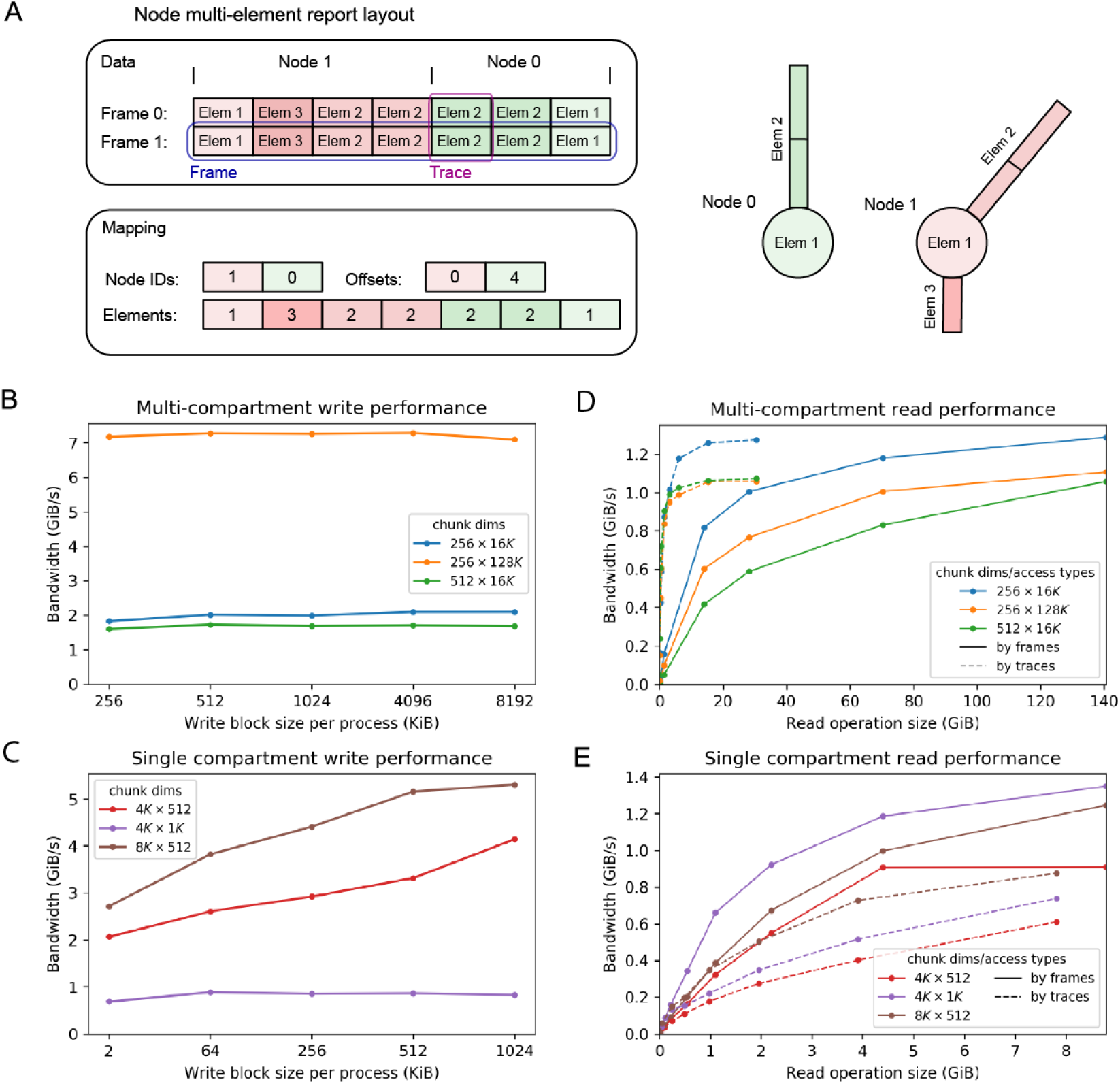
Recordings of activity in SONATA format. (A) Layout of a multi-compartment report. The dataset is a matrix where each frame (set of values at one point in time) is a row and columns represent traces (the time series of all values associated with one element). All the elements of a node are contiguous within a frame, but nodes may not appear sorted by GID. The position of the first element of each node is indicated by the offset array. Node elements can appear multiple times (e.g. morphological sections with multiple electrical compartments). (B-E) Examples of read/write performance (see **Methods**). Write performance (B, C) and read performance (D, E) of multi-compartment reports (B and D) and single compartment reports (C and E) is measured as bandwidth (amount of data written/read per time unit). Three different HDF5 chunk dimensions (specified in the legend, note that the K suffix indicates multiplication by 1024) were evaluated to demonstrate that high effective bandwidth can be obtained. In the reading evaluation, data was read by frames (continuous lines) and by traces (dotted lines) in single operations of different sizes to demonstrate the flexibility and high performance of the SONATA format; in the writing evaluation, data was only written by frames (continuous lines), which imitates the way most simulators generate data.

In the resulting design, data are stored in a single N⨉M matrix dataset, with rows being frames and columns being traces, whereas extra metadata provides a *mapping* between (cell, element) identifiers and columns within the frame (**Fig. 3A**). The format provides substantial flexibility, in particular permitting one to save different types and amounts of information for different cells. For example, one can choose to save membrane voltage and synaptic currents for all compartments and all synapses for a few cells, only somatic membrane voltage for several other cells, and nothing at all for all the other cells. This design also readily represents non-cell-element time series reports. In this case, instead of the cell elements, each row represents a channel storing a particular time series -- for example, an electrode at which the extracellular voltage is recorded.

#### Performance benchmarks

**Fig. 3B-E** illustrates the effective I/O bandwidth (amount of useful data read/written per time unit) of SONATA multi-compartment and single-compartment reports, using 26,576 neurons (41,389,269 reported cell elements) with 1,000 time steps for the former and 217,000 neurons with 130,000 time steps for the latter (see **Methods**). We considered (i) the amount of data read/written, (ii) HDF5 chunk dimensions, (iii) only for write benchmarks ― the amount of data written at each write operation (block size per process), and (iv) only for read benchmarks ― the direction in which data is accessed (by frames or by traces). We did not consider the latter option in the write benchmark because simulators typically generate data which is ordered temporally, i.e. in frames.

Note that HDF5 provides a storage layout in which the dataset is split into fixed size “chunks” (see **Methods**). Chunking is essential for obtaining good performance with arbitrary access patterns, and for that reason is supported in SONATA. However, SONATA does not prescribe specific chunking, and taking advantage of chunking to optimize read/write performance for specific applications is up to the specific software implementations that use SONATA.

The benchmarks in **Fig. 3B-E** show that SONATA supports high read and write performance. The write performance reaches several GiB/s. In the case of multi-compartment reports, the HDF5 chunk size is the main determinant of the effective write performance (**Fig. 3B**). This is due to the overhead caused when using smaller HDF5 chunk dimensions, as the increase in absolute number of HDF5 chunks makes the support data structures in the file larger. On the contrary, in single-compartment reports (**Fig. 3C**) the amount of data written by each process at each write operation affects performance, since writing data in small block sizes is not efficient. Here the performance is also affected by the fact that, in some cases, multiple processes write to the same HDF5 chunk, which leads to lower effective bandwidth (compare 4K ⨉ 512 vs 4K ⨉ 1K). The read performance tests (**Fig. 3D, E**) were run on a single-node, single-thread configuration, because this is the typical scenario of analysis and visualization use cases. In all cases, read bandwidth improves as the number of contiguous cells per operation increases and reaches 1 GiB/s and above.

### An example of a large-scale model: a network model of the layer 4 of mouse cortical area V1

To provide a realistic example of handling large-scale biologically detailed networks with SONATA, we consider the recently published network model of the layer 4 of the mouse primary visual cortex (area V1) (Arkhipov et al., 2018). The model consists of 45,000 neurons (representing more than half of layer 4 neurons in V1) and employs realistic patterns of highly recurrent connectivity. The central portion of the model (**Fig. 4A**) consists of 10,000 neurons modeled using a biophysically detailed, compartmental approach, whereas the remaining 35,000 neurons are modeled using a much simpler point-neuron, leaky integrate-and-fire (LIF) approach and serve mainly to prevent boundary artifacts. This hybrid model contains ~40 million edges for connections between explicitly modeled nodes and another ~8 million edges from ~10,000 external virtual nodes providing external spiking inputs. In the original study, the model was subjected to a battery of visual stimuli (movies), and the results were compared to published work and new in vivo experiments (Arkhipov et al., 2018) (see an example of spiking output in **Fig. 4B)**.

**Figure 4.**
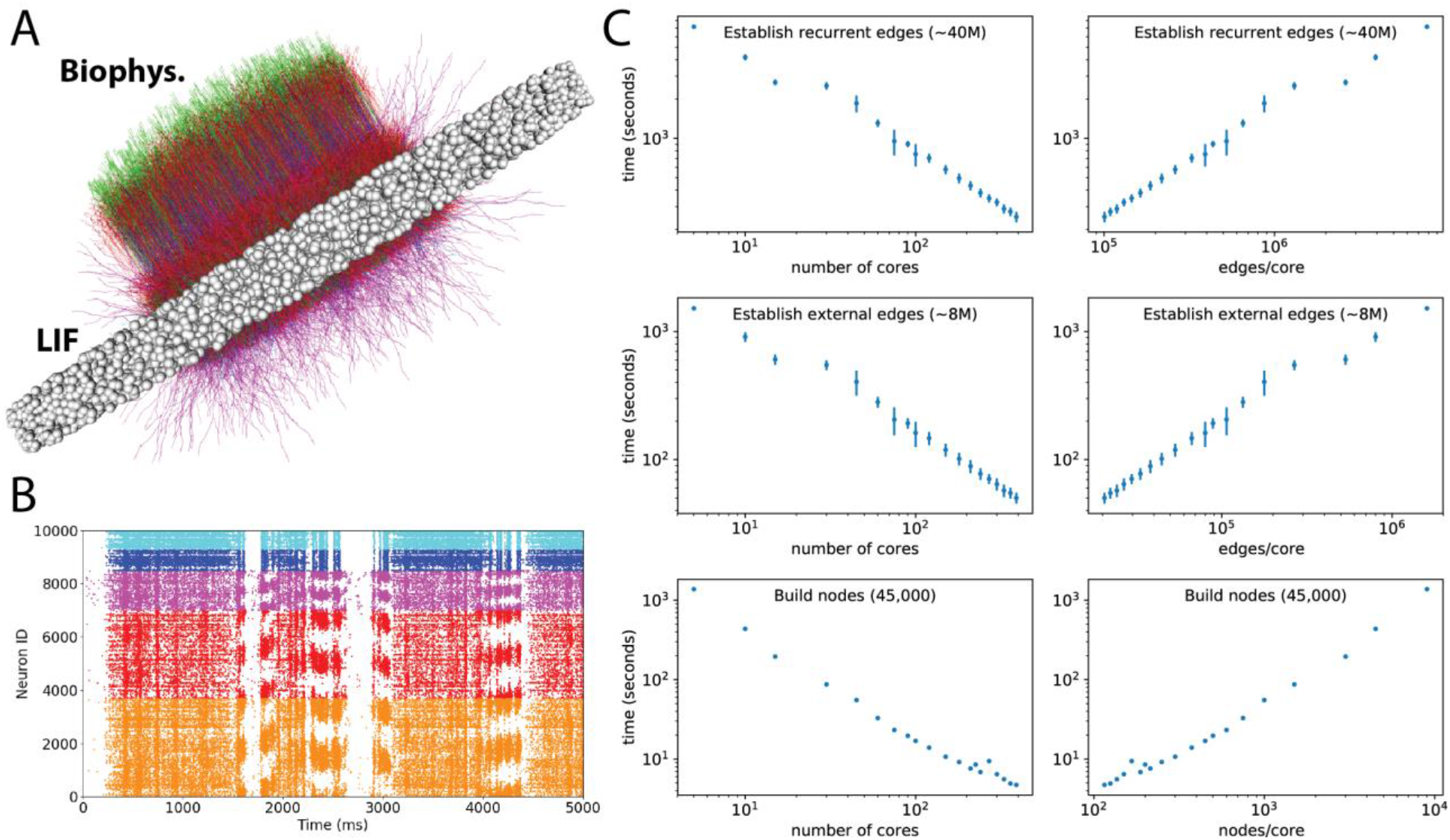
A 45,000-neuron hybrid network model of the layer 4 of mouse cortical area V1. (A) Visualization of the network model, which consists of 10,000 biophysically detailed neurons (colored morphologies) in the center and 35,000 point neurons (white balls) forming an annulus around the biophysical neurons to prevent boundary artifacts. (B) An example raster plot output from a simulation of the layer 4 model. Shown are the spikes of 10,000 biophysical neurons in response to a clip from a natural movie. Colors indicate the five types of neurons: excitatory Scnn1a (orange), Rorb (red), Nr5a1 (magenta) and inhibitory PV1 (blue) and PV2 (cyan). See details in (Arkhipov et al., 2018). (C) Benchmarks for instantiating different parts of the layer 4 model. The left and right column show the same data: against the number of CPU cores used for simulation on the left and against the number of edges or nodes per core on the right.

**Fig. 4C** shows benchmarks for loading the layer 4 model in SONATA format for simulation in NEURON (Carnevale and Hines, 2006) using the BMTK’s BioNet module (Gratiy et al., 2018), performed on cluster partitions from 5 to 390 CPU cores. The times required to build the nodes, establish edges from the external virtual nodes, and establish edges among the explicitly simulated, recurrently connected nodes are shown (note that these times include both reading the files and instantiating NEURON objects). Two views of the same data are presented: (i) scaling with the number of cores and (ii) scaling with the number of edges or nodes per core. The scaling is approximately linear (with a slope close to 1) starting at about 32 cores. The overall simulation setup time is dominated by the recurrent connections, which are about 5 times more numerous than the virtual input connections and take about 5 times longer to set up.

For a typical use case of hundreds of CPU cores, the 45,000-neuron hybrid layer 4 network model requires <10 s for instantiating nodes, <50 s for external edges, and ~4 minutes for recurrent edges, resulting in ~5-minute setup time total. Using uncompressed HDF5 files, the total size of network files, including recurrent and feedforward network connections, is ~2.4 GB (see http://portal.brain-map.org/explore/models/l4-mv1). Thus, for this considerably large and detailed model, SONATA supports modest loading times and storage space footprint. We also previously demonstrated good scaling of simulation time for this model (Gratiy et al., 2018).

### Ecosystem support

SONATA is a free format open for community development. Anyone wishing to add SONATA support to a Python based application may utilize the PySONATA Python API hosted at GitHub and developed jointly by the Allen Institute and Blue Brain Project (BBP). Multiple tools from these two organizations and other modeling and standardization initiatives already implement SONATA support (**Fig. 5**).

**Figure 5.**
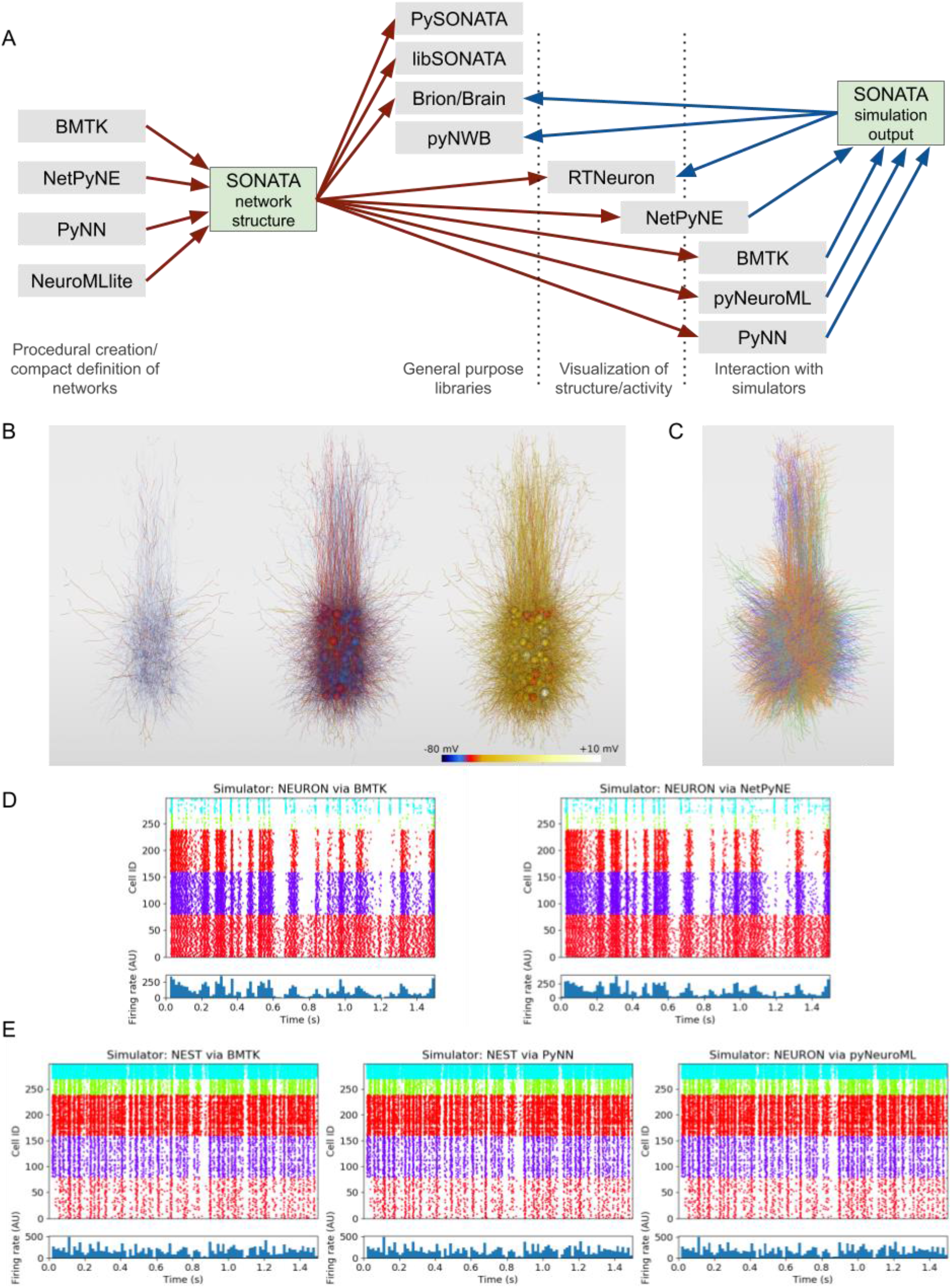
Support for SONATA in simulators and libraries. (A) Overview of applications which can generate SONATA files (containing either a description of a network structure or simulation output) and the various categories of applications which can read SONATA, including general purpose libraries, visualization tools, and simulation packages. The software packages BMTK, NetPyNE, PyNN, and pyNeuroML can read SONATA network descriptions for execution in the simulation engines NEURON and NEST. The pyNWB package provides a programming interface for reading and writing neurophysiology data (either from experiments or from simulations) in the NWB:N 2.0 format. (B) RTNeuron visualization. Sample renderings at 3 simulation timesteps of an example network with 300 biophysically detailed cells, with somatic and dendritic compartments colored according to the simulated membrane potential. The biophysical 300-cell network, as well as its point-neuron counterpart, were created via the model-building scripting interface in BMTK and saved using SONATA. These two models are used in all subsequent panels here. (C) Rendering of the same model as in (B) using the NetPyNE GUI. Each cell is colored according to which of the 5 node types it belongs. (D) The 300-cell biophysically detailed example from (B) and (C) simulated in NEURON using BMTK (left) and NetPyNE (right). (E) A network with 300 integrate and fire neurons generated by BMTK, and simulated in NEST via BMTK (left), NEST after importing the SONATA files into PyNN (middle) and NEURON after conversion of the network to NeuroML by pyNeuroML. Each raster plot in (D) and (E) is accompanied by a panel underneath showing population firing rate (arbitrary units).

Below we briefly describe examples of using these tools to construct, read, write, visualize, and simulate network models in SONATA format. Note that, in general, when different simulators load one SONATA model for simulation, bitwise agreement between their outputs is not guaranteed. The reasons for that are non-standardized processing of certain data in simulation software packages, different approaches for instantiating initial conditions, etc. For a real-life example, consider that loading SWC morphologies in NEURON can be done using different functions (e.g., hoc or Python), which employ different numerical precisions; as a result, simulation outputs will not be bitwise identical, but will be only statistically the same to the level permitted by the precision discrepancy in morphologies. Nevertheless, SONATA constrains a vast variety of important degrees of freedom in network simulations, enabling statistically similar results between simulators and bitwise reproducibility within a simulator with fixed software code.

Although SONATA has been originally developed to support very large and biologically complex simulations, it is fully consistent with more typical smaller-scale and less complex applications. For example, it is rather common for modelers to use conceptual rules implemented in a few lines of code to generate model geometries and connections. These approaches are fully supported by BMTK, Brion/Brain, pyNeuroML, PyNN, and NetPyNE described below -- in addition to the advanced capabilities of these tools to build and carry out very sophisticated, data-driven, large-scale network simulations. Each of these software packages can generate models using such high-level conceptual definitions, and in fact the examples illustrated in **Fig. 5** were generated in such a simple way using the BMTK’s model building module. The important new contribution that SONATA makes is a standardized, efficient format for exchanging generated network structures, as well as simulation results, between these applications. That is showcased in **Fig. 5**, where the BMTK-generated models are simulated using several other tools. Furthermore, it is important to note that large scale biologically realistic models (e.g., (Arkhipov et al., 2018; Markram et al., 2015)) often require as much or even more time to build than to run a single simulation, and then saving model instantiations becomes very important, whereas for small models this may be simply unnecessary. However, for sharing models with the community, and especially across simulator platforms, the ability to save all instantiated parameters of models and simulations systematically -- as provided by SONATA -- becomes important for large and small models alike. The examples in **Fig. 5** are relatively small, 300-neuron models, illustrating use cases that are more common in the field than the very large simulations with tens of thousands of neurons (**Fig. 4**).

Currently, SONATA is not natively supported by the widely used simulation engines NEURON and NEST, but the tools described below provide convenient interfaces to NEURON and NEST and enable simulations with SONATA using these two engines. In the future, implementation of native support in NEURON and NEST could be useful for systematic conversion of older, existing models (which are typically stored as software code) to SONATA format by instantiating these models in NEURON or NEST environment from the original code and then saving as SONATA files.

#### PySONATA

PySONATA is a Python based API for reading SONATA files, open-sourced under a BSD license and maintained as an official tool of the SONATA working group (https://github.com/AllenInstitute/sonata). Users wishing to begin integrating the SONATA format into their own software are encouraged to use the PySONATA Python modules.

Examples of how to use the module can be found at https://github.com/AllenInstitute/sonata/blob/master/src/pysonata/docs/Tutorial%20-%20pySONATA.ipynb.

#### The Brain Modeling Toolkit

The Brain Modeling Toolkit (BMTK; https://github.com/AllenInstitute/bmtk) is a Python based package for building, simulating and analyzing large-scale neural networks across different levels of resolution. The BMTK is open-sourced under a BSD-3 license and has full support for generating and reading the SONATA data format (**Fig. 5**). Modelers can use the BMTK Builder submodule to create their own SONATA based networks from scratch. It supports cell template files, electrophysiological parameters, and morphology from the Allen Cell Types Database (http://celltypes.brain-map.org/) (Gouwens et al., 2018b; Teeter et al., 2018) as well as other cell model formats, including NeuroML2 (Cannon et al., 2014; Gleeson et al., 2010), NEURON hoc files (Carnevale and Hines, 2006), or even user defined Python functions. For simulations, BMTK relies on an increasing array of simulation engines (NEURON (Carnevale and Hines, 2006), NEST (Gewaltig and Diesmann, 2007), Dipde (Cain et al., 2016), etc.), which allow users to run simulations of SONATA networks using either multi-compartment, point, or population based representations. The results of these simulations, regardless of the underlying simulator used to run them, are transformed into SONATA output format, allowing networks built and run with BMTK to be analyzed and visualized by any third-party software that supports SONATA. **Fig. 5B** and **5C** show a network with 300 biophysically detailed cells, in SONATA format, generated using BMTK and visualized with RTNeuron and NetPyNE, respectively. The results of simulations of this network using BMTK and NetPyNE are shown in **Fig. 5D**. **Fig 5E** shows simulations of a network of 300 integrate and fire neurons created with BMTK and simulated using BMTK, PyNN, and pyNeuroML.

#### Brion/Brain

The Blue Brain’s C++ libraries for handling large scale data and simulation setup, Brion/Brain (https://github.com/BlueBrain/Brion), provide partial support for SONATA. Currently Brion provides a low level API to read circuit and simulation JSON configurations, spike and multi-compartment simulation outputs, SWC morphologies and query nodes in HDF5 files. It also provides a single threaded writer for multi-compartment simulation output reports. Brain provides a higher level API that makes it easier to work with full networks. All this functionality is also available in Python through the associated Python wrapping module.

#### libSONATA

Blue Brain’s libSONATA (https://github.com/BlueBrain/libsonata) is a library that provides support to read SONATA files. The library is open-sourced under a LGPLv3 license and offers an API for both Python and C++ applications. Currently libSONATA supports reading circuit files, including nodes and edges populations.

#### RTNeuron

Blue Brain’s RTNeuron (Hernando et al., 2013) is a framework for visualizing detailed neuronal network models and simulations. As it relies on Brion/Brain for data access, it currently provides basic support to visualize SONATA circuits and simulations. For instance, **Fig. 5B** illustrates the RTNeuron visualization of a model of 300 biophysically detailed neurons, provided as an example in the SONATA specification GitHub repository (https://github.com/AllenInstitute/sonata/tree/master/examples/300_cells). Here, one can see neuronal morphologies and the distribution of membrane voltage across the electrical compartments comprising these morphologies as the simulation evolves over time.

#### pyNeuroML

NeuroML (Cannon et al., 2014; Gleeson et al., 2010) is a standardized format based on XML for declaratively describing models of neurons and networks in computational neuroscience. Cellular models which can be described range from simple point neurons (e.g. leaky integrate and fire) to multicompartmental neuron models with multiple active conductances. Networks of these cells can be specified, detailing the 3D positions or populations, connectivity between them and stimulus applied to drive the network activity.

Multiple libraries have been created to support user adoption of the NeuroML language, including jNeuroML (https://github.com/NeuroML/jNeuroML) in the Java language and pyNeuroML (https://github.com/NeuroML/pyNeuroML) in Python. The latter package also gives access to all of the functionality of jNeuroML (including the ability to convert NeuroML to simulator specific code, e.g. for NEURON) through Python scripts, by bundling a binary copy of the library. PyNeuroML has recently added support for importing networks and simulations specified in the SONATA format and converting them to NeuroML. A related package currently under development, NeuroMlite (https://github.com/NeuroML/NeuroMLlite) allows compact description of networks and can export the generated structures to SONATA. **Fig. 5E** shows a simulation of 300 integrate and fire cells in SONATA which has been imported by pyNeuroML, converted to NeuroML and executed in the NEURON simulator.

#### PyNN

PyNN is a simulator-agnostic Python API for describing network models of point neurons, and simulation experiments with such models (Davison et al. 2009). A reference implementation of the API for the NEURON, NEST and Brian simulators is available (http://neuralensemble.org/PyNN), and a number of other simulation tools, including neuromorphic hardware systems, have implemented the API (Brüderle et al., 2011; Rhodes et al., 2018). PyNN models can be converted to and from the NeuroML and SONATA formats with a single function call. **Fig. 5E** illustrates an example where a model in SONATA format was loaded using the PyNN “serialization” module, a simulation was carried out using the PyNN NEST backend, and simulation output was saved in the SONATA format.

#### NetPyNE

NetPyNE (www.netpyne.org; (Dura-Bernal et al., 2019)) is a package developed in Python and building on the NEURON simulator (Carnevale and Hines, 2006). It provides both programmatic and graphical interfaces that facilitate the definition, parallel simulation, and analysis of data-driven multiscale models. Users can provide specifications at a high level via its standardized declarative language. NetPyNE supports both point neurons and biophysically-detailed multi-compartment neurons, as well as NEURON’s Reaction-Diffusion (RxD) molecular-level descriptions. The tool includes built-in functions to visualize and analyze the model, including connectivity matrices, voltage traces, raster plot, local field potential (LFP) plots and information transfer measures. Additionally, it facilitates parameter exploration and optimization by automating the submission of batch parallel simulation on multicore machines and supercomputers.

NetPyNE network model instantiations can be converted to and from the NeuroML and SONATA formats. SONATA complements NetPyNE by providing a standardized and efficient format to store and exchange large network models. This enables using other simulation tools to run and explore models developed with NetPyNE, and vice versa. As an example, we imported the 300-cell SONATA example with multicompartment cells into NetPyNE, visualized it using the NetPyNE GUI (**Fig. 5C),** and carried out a NetPyNE simulation (**Fig. 5D).**

#### Neurodata Without Borders: Neurophysiology 2.0

Neurodata Without Borders: Neurophysiology (NWB:N) 2.0 is a data format for standardizing experimental data across systems neuroscience. We developed an extension for NWB:N 2.0 to accommodate large-scale simulation data, and developed a conversion script from SONATA to NWB:N 2.0 (https://github.com/ben-dichter-consulting/ndx-simulation-output) (**Fig. 5A**). This allows stimulated data to be stored side-by-side with experimental data and facilities comparative analysis between simulation and electrophysiology or calcium imaging experiments.

## Discussion

We have described SONATA, an open-source data format developed to answer the challenges of modern computational neuroscience, especially those inherent in large-scale data-driven modeling of brain networks. It is designed for memory and computational efficiency, as well as for working across multiple platforms, and at the same time enabling as much flexibility as possible for diverse applications. To achieve this, SONATA relies on commonly used data formats such as CSV, HDF5, and JSON, which can be used across platforms, can be read and written by many existing libraries in various programming languages, and (especially in the case of HDF5) have been proven to work efficiently in parallel computations with very large datasets. The SONATA specifications include network descriptions, simulation configuration, and input or output activity. Close cooperation with existing standardisation and simulator independent specification initiatives like NeuroML, PyNN, and NWB:N has helped to increase synergy with existing formats, and has ensured compatibility with languages and tools already in use in the community.

The flexibility of the SONATA specification is ensured by several design criteria. First, the design leaves it up to users to decide which attributes are shared within node or edge type vs. which are unique to specific nodes or edges. Second, it allows limitless creation of user-defined attributes and maintains only a small number of reserved fields. And third, via a hierarchy of types, populations, and groups of nodes/edges, it permits specification of hybrid models that may include biophysically detailed neurons, point neurons, and many other model types, all in one network model.

While SONATA offers computationally efficient solutions for storing many model properties, we did not attempt to reinvent file formats for all properties. For example, SONATA utilizes the well established ASCII-based SWC format for neuronal morphologies. We did not develop a computationally optimized binary format for morphologies because their footprint in terms of storage or computational demand is typically small. In the case of the Layer 4 model (**Fig. 4**), loading SWC morphologies takes ~60% of the time of building nodes, but that expense is dwarfed by the time it takes to establish connections (~300 s for external and recurrent connections vs. ~5 s for nodes). Thus, we opted to develop efficient binary solutions only for computationally demanding model properties, otherwise relying on widely used formats such as SWC.

The SONATA community and ecosystem include multiple groups with diverse interests and are growing due to the open-source design. Initially developed jointly by the Allen Institute and the Blue Brain Project, SONATA is now supported by tools from many teams. As described above, tools such as BMTK (Gratiy et al., 2018), RTNeuron (Hernando et al., 2013), PyNN (Davison et al., 2009), NeuroML (Cannon et al., 2014; Gleeson et al., 2010), and NetPyNE (Dura-Bernal et al., 2019) include SONATA support. Functionality for conversion between SONATA and NWB:N (Ruebel et al., 2019) also exists. The SONATA data format and framework are reflected in the free and open-source PySONATA project hosted on GitHub (https://github.com/AllenInstitute/sonata), which is intended as a key resource for those wishing to add support for SONATA to their applications and includes specification documentation, open-source reference application programming interfaces, and model and simulation output examples.

As an open living format, SONATA may be extended in the future to reflect developments in modeling and in experimental neuroscience. In turn, we invite experimentalist colleagues to explore SONATA’s applicability to their circumstances, as the SONATA framework provides an efficient description for a variety of network properties. Such cross-pollination will help improve reproducibility and facilitate collaboration between experimental and computational neuroscientists.

## Methods

### JSON, CSV, and HDF5

#### JSON

JSON (JavaScript Object Notation) is a data exchange format that is easy for both humans and machines to read and write. Being text based, JSON is platform and language independent. Data organization is based on two common structures: key-value pairs and ordered lists, which have equivalents in almost all programming languages.

#### CSV

CSV stands for “*comma-separated values”* and it is a very common way of laying out tabular data in text files. CSV is not a standard *per se*; the choices that have been made for SONATA are described in the official specification. It should be noted that, although the CSV abbreviation suggests comma as a separator, CSV files can use many types of separator, and, in fact, SONATA format specifies spaces as preferred separators for CSV.

#### HDF5

HDF5 (Hierarchical Data Format version 5) is a technology designed for storing very large heterogeneous data collections and their metadata in a single container file. HDF5 defines a binary container file format for which the HDF Group provides an implementation in C. Bindings for several other languages exist as well. Basic concepts of HDF5 include groups, datasets and attributes. Making an analogy to filesystems, groups are similar to directories and datasets to files. The main differences between HDF5 and a general purpose filesystem are that a) a dataset is not a stream of bytes like a file, but consists of a multidimensional array with a single data type for all values and that b) groups and datasets can be annotated by means of attributes. HDF5 defines some basic data types common to most programming languages: integers, floats, strings. Data can be stored linearly (the elements of a dataset are stored in increasing order, according to their index and dimension) or in “chunks” for computational efficiency (the order in how dataset elements is interleaved according to their index and dimension; for details, see https://support.hdfgroup.org/HDF5/doc/Advanced/Chunking/).

### Benchmarking

#### Edge file benchmarks

The performance of navigating through an edge file in SONATA format is illustrated in **Fig. 6**, which shows the results of selecting 1000 neurons and accessing one arbitrary property of all the edges of the selected neurons in the 45,000-cell recurrently connected model of Layer 4 of mouse V1(Arkhipov et al., 2018) On average each cell receives input from 438.8 neighbors with the number and strength of synapses between any two cells being determined by source and target cell types. The network file contains over 39.2 million unique synapses partitioned into two groups, those synapses that target multicompartment neurons and those that target point points. Connections that target point neurons only require synaptic strength variable, while those that target multi-compartment neurons also require information about section number and segment distance for each synapse. The HDF5 edge file is 1.9 GB in size.

**Figure 6.**
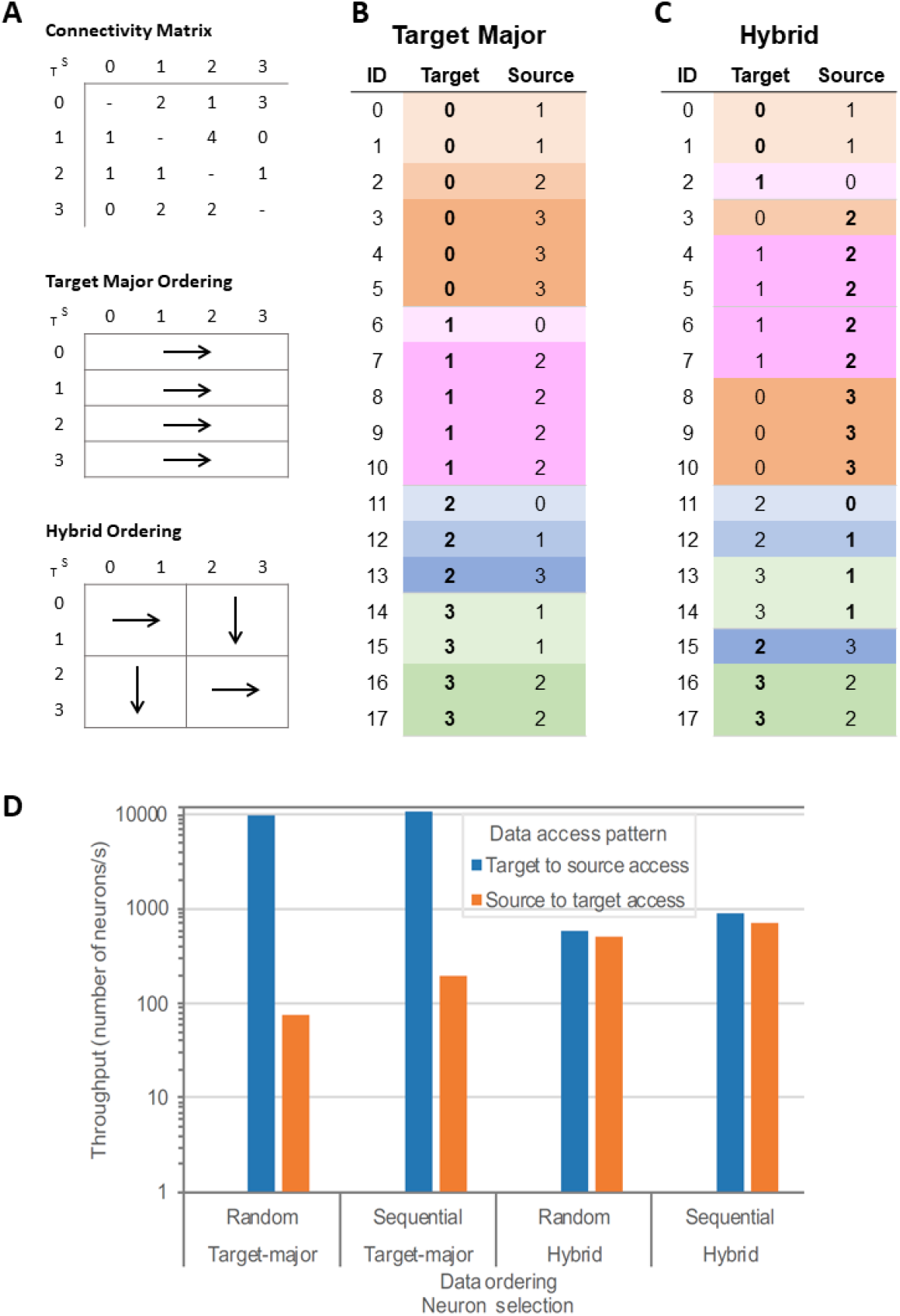
Target major and hybrid ordering of edges. (A) A simple example of connectivity matrix (the number within each matrix element indicates the number of edges -- i.e., synapses --between the two nodes) and schematics of target major and hybrid orderings. (B) and (C) Edge lists representing edges from the connectivity matrix in (A), sorted according to target major (B) or hybrid (C) ordering. (D) Throughput of accessing edge information for target major or hybrid ordering of edges in the SONATA files in a 45,000-cell model of Layer 4 of mouse V1 (Arkhipov et al., 2018). The target-to-source and source-to-target access patterns are illustrated with either random or sequential selection of target or source neurons.

The benchmarks were conducted on an HPE SGI 8600 supercomputer. Each compute node had two Intel Xeon Gold 6140 CPUs (each with 18 cores at 2.30 GHz) and 768 GB of DRAM. Nodes were connected through a Mellanox Infiniband (IB) EDR fabric to two GS14K storage racks with a total storage capacity of 4 PB. The computing system was running Linux 3.10.0 and the filesystem was GPFS 4.2.3-6, configured with 4 MiB block size. The storage system did not have dedicated metadata drivers. The software components used and their versions are the following: glibc 2.25-49, gcc 6.4, boost 1.58, HDF5 1.10.1, Python 2.7, numpy 1.13.3 and MPI 2.16 provided by HPE.

For reference, the maximum average read bandwidth obtained in pure I/O benchmark experiments with IOR (https://ior.readthedocs.io/en/latest/) in this machine is 5.6 GiB/s using 1 single core accessing a 1 GiB file in 4 MiB blocks. The maximum average write bandwidth measured is 9.5 GiB/s using 8 cores from 1 node writing 1 GiB per core in 4 MiB operations to a shared file. POSIX I/O was used to obtain both measurements.

To illustrate SONATA’s performance and flexibility, we use examples of ordering the edges data in two different ways (**Fig. 6A)**: target-major (**Fig. 6B**), where data is sorted according to the ID of the target neuron (increasing), and hybrid ordering (**Fig. 6C**), where the connectivity matrix is divided in blocks, and edges inside each block are enumerated, alternating (from block to block) between source-major and target major orderings. We also compare the impact of selecting 1000 neurons randomly or sequentially.

Note that SONATA supports arbitrary ordering of edges, and the two variants tested in the benchmarks are only for demonstration purposes.

A target-major sort is more efficient for instance in the case of a simulator creating the synapses on the target cell when instantiating the network. A source-major sort (data sorted according to the ID of the source neuron increasing) is favorable to analysis of efferent connectivity of large network. The hybrid ordering is a compromise between the target-major and source-major ordering.

**Fig. 6D** shows that ordering has an impact on the performance of data access (whereas selecting neurons randomly or sequentially does not impact performance substantially). By using target-major ordering (or its symmetric source-major ordering) one can achieve optimal performance when accessing data in the same access pattern as the ordering, but accessing data in the opposite direction is much less efficient, by a factor of ~100. Ordering data in a hybrid manner is a compromise to get balanced performance between the source-to-target and target-to-source access patterns, but in this case the performance is not as good as the optimal performance for non-hybrid ordering. Due to such large discrepancies, the SONATA format specification leaves the choice of ordering open to users. Note that source-target pairs for each edge are always defined in the edge files in the same way; it is the indexing of these edges that may differ depending on user requirements. This means that the edges can always be read, but reading speed for a particular application will depend on the choice of indexing, and this choice should be made based on the desired application. Examples in **Fig. 6D** indicate that a rather high performance can be achieved (close to 10,000 neurons processed per second for their edge attributes) in optimal cases, but users should take advantage of the flexibility of SONATA specification to use edge ordering that is most suitable for their needs. In situations where high performance for various access patterns is essential, solutions may include two or more copies of edge files with different orderings for different use cases.

#### Simulation output benchmarks

The simulation output benchmarks (**Fig. 3**) were run on the aforementioned HPE SGI 8600 system. Since most simulators can run in parallel (multi-thread and/or multi-process), the benchmarking of the report generation was also done in parallel, on 16 nodes and 36 processes per node (using 1 core per process). All processes were periodically dumping data to a single, shared HDF5 file in the SONATA format. At each write operation, each process was writing several columns at its designated frame/trace region. The amount of data written at each operation is presented as the “Write block size per process” illustrated in the performance plots (the write block size applies for each process and for each write operation).

Write benchmarks made use of the Neuromapp library (https://github.com/BlueBrain/neuromapp, revision f03d3ea) (Ewart et al., 2017), which uses parallel HDF5 and MPI underneath. Read benchmarks were implemented using the Python binding of Brion/Brain (revision c16a694), the testing and plotting code can be found in the SONATA github repository in the benchmarks branch.

#### Loading of simulation data

Benchmarks for loading simulation data (**Fig. 4C**) were obtained for the full simulation of the 45,000-neuron recurrently connected model of Layer 4 of mouse V1 (Arkhipov et al., 2018). Figure **Fig. 4C** shows the amount of time required to parse through the SONATA network files and instantiate the in-memory cell and synaptic objects to run a full NEURON (Carnevale and Hines, 2006) simulation. Each simulation was instantiated with a computing cluster of Intel Xeon E5 processors (each core either 2.1 or 2.2 GHz), using a minimum of 5 cores and a maximum of 390 cores. The network was built using the Brain Modeling Toolkit with Python 3.6 and NEURON 7.5 with Python bindings.

## Acknowledgements

This project/research has received funding from the European Union’s Horizon 2020 Framework Programme for Research and Innovation under the Specific Grant Agreement No. 785907 (Human Brain Project SGA2) and from the EPFL Blue Brain Project (funded by the Swiss government’s ETH Board of the Swiss Federal Institutes of Technology). SD-B was funded by NIH grants U01EB017695, R01EB022903 and 2R01DC012947-06A1, and New York State grant DOH01-C32250GG-3450000. PG was supported by the Wellcome Trust (212941, 101445). This work benefited from interactions between the authors as part of the International Neuroinformatics Coordinating Facility (INCF) Special Interest Group on Standardised Representations of Network Structures. We are grateful to Christof Koch for inspiring and supporting this work and to Michael Hines for many helpful discussions and suggestions. We wish to thank the Allen Institute founder, Paul G. Allen, for his vision, encouragement, and support.

